# Uncovering the repertoire of endogenous flaviviral elements in *Aedes* mosquito genomes

**DOI:** 10.1101/124032

**Authors:** Yasutsugu Suzuki, Lionel Frangeul, Laura B. Dickson, Hervé Blanc, Yann Verdier, Joelle Vinh, Louis Lambrechts, Maria-Carla Saleh

## Abstract

Endogenous viral elements derived from non-retroviral RNA viruses were described in various animal genomes. Whether they have a biological function such as host immune protection against related viruses is a field of intense study. Here, we investigated the repertoire of endogenous flaviviral elements (EFVEs) in *Aedes* mosquitoes, the vectors of arboviruses such as dengue and chikungunya viruses. Previous studies identified three EFVEs from *Ae. albopictus* and one from *Ae. aegypti* cell lines. However, in-depth characterization of EFVEs in wild-type mosquito populations and individuals *in vivo* has not been performed. We detected the full-length DNA sequence of the previously described EFVEs and their respective transcripts in several *Ae. albopictus* and *Ae. aegypti* populations from geographically distinct areas. However, EFVE-derived proteins were not detected by mass spectrometry. Using deep sequencing, we detected the production of piRNA-like small RNAs in antisense orientation, targeting the EFVEs and their flanking regions *in vivo*. The EFVEs were integrated in repetitive regions of the mosquito genomes, and their flanking sequences varied among mosquito populations from different geographical regions. We bioinformatically predicted several new EFVEs from a Vietnamese *Ae. albopictus* population and observed variation in the occurrence of those elements among mosquito populations. Phylogenetic analysis of an *Ae. aegypti* EFVE suggested that it integrated prior to the global expansion of the species and subsequently diverged among and within populations. Together, this study revealed substantial structural and nucleotide diversity of flaviviral integrations in *Aedes* genomes. Unraveling this diversity will help to elucidate the potential biological function of these EFVEs.

**Importance:** Endogenous viral elements (EVEs) are whole or partial viral sequences integrated in host genomes. Interestingly, some EVEs have important functions for host fitness and antiviral defense. Because mosquitoes also have EVEs in their genomes, we decided to thoroughly characterized them to lay the foundation of the potential use of these EVEs to manipulate the mosquito antiviral response. Here, we focused on EVEs related to the *Flavivirus* genus, to which dengue and Zika viruses belong, in *Aedes* mosquito individuals from geographically distinct areas. We showed the existence *in vivo* of flaviviral EVEs previously identified in mosquito cell lines and we detected new ones. We showed that EVEs have evolved differently in each mosquito population. They produced transcripts and small RNAs, but not proteins, suggesting a function at the RNA level. Our study uncovers the diverse repertoire of flaviviral EVEs in *Aedes* mosquito populations and suggests a role in the host antiviral system.

## Introduction

Endogenous viral elements (EVEs), also known as viral fossils, are whole or partial viral sequences integrated in host genomes (1). When viral DNA integration occurs in the germline, it can be inherited and retained in the host genome as evidence of ancient viral infections old from millions of years. Retrovirus-derived EVEs are the best-known examples since retroviruses actively integrate their DNA into the host genome as part of their life cycle during infection. However, single-stranded DNA virus-derived elements were detected in plants (2) and more recently, non-retrovirus-derived EVEs have been shown in various animal hosts (3-7). Indeed, recent advances in bioinformatics have dramatically changed the landscape of paleovirology. *In silico* surveys were used to screen for EVEs in various animal genomes and identified a number of non-retroviral EVEs sequences belonging to several virus families (5-8). The integrated viral elements mostly accumulate random mutations that render them inactive. In several instances, however, EVEs have maintained open reading frames (ORFs) and produce functional proteins that can serve during infections by closely related viruses (9-12). For example, endogenous bornavirus-like nucleoproteins from thirteen-lined ground squirrel, *Ictidomys tridecemlineatus*, (itEBLN) is the first non-retroviral EVE demonstrated to serve as a negative regulator against infection by a related virus (10). Overexpression of itEBLN inhibits borna disease virus (BDV) infection in mammalian cell lines, presumably by decreasing BDV polymerase activity. More recently, a virophage mavirus, which parasites *Cafeteria roenbergensis* virus (CroV), was shown to be endogenized as an EVE in a marine flagellate, *Cafeteria roenbergensis* (*C. roenbergensis*) (9). CroV infection activates the endogenized mavirus genes in *C. roenbergensis* and produces infectious mavirus particles. These particles are secreted and protect surrounding flagellates from subsequent CroV infection. These studies demonstrate that EVEs can play critical roles in the host antiviral system.

Like most species examined, *Aedes* mosquitoes also have EVEs in their genomes. *Aedes* (*Ae*.) *aegypti* and *Ae. albopictus* are major vectors of arthropod-borne viruses (arboviruses) such as dengue virus (DENV) and Zika virus (ZIKV). DENV and ZIKV are members of the *Flavivirus* genus, which consists of enveloped viruses with a positive-sense, single-stranded RNA genome. In addition to these medically important mosquito-borne flaviviruses, *Aedes* mosquitoes in nature are infected by insect-specific flaviviruses (ISFs) such as cell-fusing agent virus (CFAV), Kamiti River virus (KRV) and *Aedes* flavivirus (AEFV) (13-15). Using *Ae. aegypti* and *albopictus* cell lines, Crochu *et al*. detected endogenous flaviviral elements (EFVEs) which showed approximately 60%, 70% and 80% amino-acid similarity to CFAV, KRV and AEFV sequences, respectively (4). In recent years, the availability of reference genome sequences for both *Ae. aegypti* and *Ae. albopictus*, and *in silico* studies confirmed the existence of various EFVEs in *Aedes* mosquitoes (5, 8). However, very limited experimental validation has been performed *in vivo*. To support our hypothesis that EVEs could be used to manipulate the mosquito antiviral response to stop arbovirus transmission to the human host, we conducted a comprehensive characterization of EVEs in *Aedes* mosquitoes representative of natural populations worldwide.

We investigated EFVEs found in two species of *Aedes* mosquitoes, *Ae. albopictus* and *Ae. aegypti* using several populations of each species sampled from geographically distinct locations. We showed the presence of EFVE DNA and RNA transcripts *in vivo*. We confirmed the production of small RNAs derived from EFVEs RNA *in vivo*. We further performed genomic DNA sequencing analysis in *Ae. albopictus* and applied an *in silico* screening procedure that identified several new EFVEs in this species. Together, our results demonstrate the ubiquitous presence of diverse EFVEs *in vivo* and contribute to understand the putative role of these elements in the antiviral defense system of mosquitoes during EFVE-related viral infection.

## Results

### Detection of endogenous flaviviral elements in *Aedes* mosquitoes

Several EFVEs in *Aedes* mosquitoes have been previously described (4, 5, 8). We focused on annotated EFVEs containing flaviviral non-structural (NS) genes, three in *Ae. albopictus* (one named CSA1, and two unnamed) and one in *Ae. aegypti* (named CSA2) (4). For simplicity, we renamed these EFVEs as: *Albopictus* Flaviviral Element (ALFE) 1 to 3, and *Aegypti* Flaviviral Element (AEFE) 1 (Fig. 1A). ALFE1-3 were originally identified in C6/36 (Ae. *albopictus*) and AEFE1 in A20 (Ae. *aegypti*) cell lines. Laboratory and field-collected *Ae. albopictus* and *Ae. aegypti* mosquitoes were positive by PCR for the NS3 and/or NS5 regions of both ALFE and AEFE (4). However, the full-length elements were not confirmed in individual mosquitoes. Because geographical origin is expected to be associated with genetic divergence due to selective pressures particular to each geographical region and/or genetic drift, we explored EFVEs in mosquitoes from different parts of the world. We used *Ae. albopictus* from Gabon and Vietnam, and *Ae. aegypti* from Cameroon, French Guiana and Thailand. First, we attempted to detect full-length DNA for ALFE1-3 and AEFE1 *in vivo*. Cell lines corresponding to each mosquito species (Aag2 for *Ae. aegypti* and C6/36 and U4.4 for *Ae. albopictus*) were used in addition to mosquito individuals. PCR showed amplicons with the expected sizes for ALFE1-3 and AEFE1 except for ALFE1 in the *Ae. albopictus* Vietnam strain (no amplification band) and ALFE2 in U4.4 cell line (a larger band) (Fig. 1B, white arrowhead). We further characterized ALFE1 in the Vietnam strain by PCR with different primer pairs. ALFE1 was amplified with a primer pair targeting the element without the first 200 base pairs (bp) at its 5’ end (Fig. 1B). This result suggests that ALFE1 in *Ae. albopictus* Vietnam differs from ALFE1 in *Ae. albopictus* Gabon, C6/36 and U4.4 cells in the first 200 bp of the element. For ALFE2, sequencing of the PCR product in U4.4 cells showed a partial ALFE2 sequence fused with unannotated mosquito sequences, suggesting that ALFE2 is recombined in U4.4 cells.

**Fig 1.**
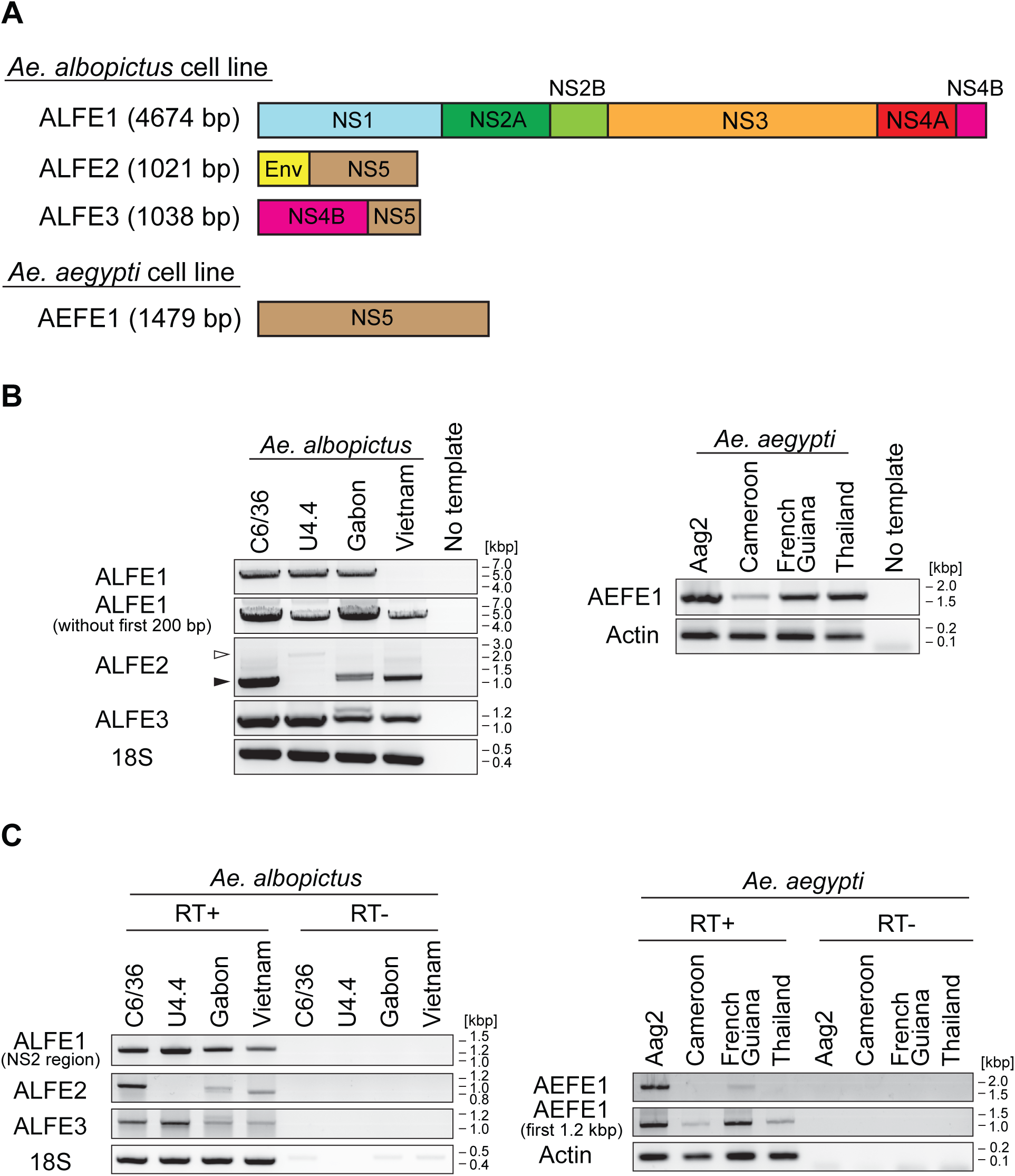
Detection of ALFE and AEFE DNA and mRNA in *Aedes* mosquitoes. (A) Schematic representation of ALFE1-3 and AEFE1. (B) Detection of ALFE1-3 and AEFE1 DNA by PCR and (C) mRNA by RT-PCR in C6/36 and U4.4 cell lines, *Ae. albopictus* Gabon and Vietnam strains (left panels) and Aag2 cell line, *Ae. aegypti* Cameroon, French Guiana and Thailand strains (right panels). Black and white arrowheads in (B) indicate ALFE2-derived bands in C6/36 cell line, Gabon and Vietnam strains, and ALFE2 fused with host sequence in U4.4 cell line. 18S and Actin genes were used as controls *for Ae. albopictus* and *Ae. aegypti*, respectively.

Next, we examined mRNA expression for ALFE1-3 and AEFE1 by RT-PCR. Previous work by Crochu *et al*. showed ALFE1 mRNA in C6/36 but not *in vivo* while ALFE2, -3 and AEFE1 mRNAs had not been tested *in vitro* nor *in vivo* (4). ALFE1 mRNA was observed with a primer pair targeting the NS2 region but not the full-length element (Fig. 1C). ALFE2 and -3 mRNAs were present in all samples except for ALFE2 in U4.4 cells. Full-length AEFE1 mRNA was observed in Aag2 cells and in the *Ae. aegypti* French Guiana strain but not in the *Ae. aegypti* Cameroon and Thailand strains. The first 1.2 kb of AEFE1 mRNA were detected in all *Ae. aegypti* samples (Fig. 1C). The *Ae. aegypti* French Guiana strain was the only to show the full-length transcript of AEFE1 among all *Ae. aegypti* strains tested. Altogether, our results indicated that ALFE1-3 and AEFE1 were established by ancient viral infections in nature and not by artificial recombination in cell culture. In addition, ALFE1-3 and AEFE1 are well conserved and their mRNAs are expressed among *Aedes* mosquito populations from different parts of the world.

### ALFE and AEFE produce piRNA-like molecules in *Aedes* mosquitoes

We previously reported that *Ae. albopictus* and *Ae. aegypti* infected with chikungunya virus (CHIKV) produce viral cDNA through endogenous retrotransposon activity and that this viral cDNA generates small interfering RNAs (siRNAs) and PIWI-interacting RNAs (piRNA) mediating viral persistence (16). CHIKV is an alphavirus from the *Togaviridae* family, which consists of enveloped viruses with a single-stranded, positive-sense RNA genome. CHIKV is also a mosquito-borne virus with great impact on human health (17). To check if ALFE1-3 and AEFE1 transcripts were also capable of generating small RNAs, as CHIKV-derived viral DNA, we re-analyzed small RNA libraries from *Ae. albopictus and Ae. aegypti*, infected or uninfected with CHIKV, that were already available in the laboratory (Fig. 2 and Fig. S1). The majority of small RNAs that mapped to ALFE1-3 and AEFE1 were 27-29 bases in length with an enrichment of uridine at the first position of the small RNA, and thus identified as primary piRNA-like molecules (Fig. 2A and Fig. S1A). We observed that piRNAs derived from these endogenous viral elements were only in antisense orientation. CHIKV infection did not affect the size distribution and profiles of the small RNAs derived from ALFE1-3 and AEFE1 (Fig. 2B and Fig. S1B). We examined the production of piRNAs targeting the flanking regions of ALFE1-3 determined by Crochu *et al* (4). We only detected production of antisense strand piRNA-like molecules on these regions, similar to ALFE1-3 (Fig. S2). In addition, the flanking regions of ALFE1 and 2, produced 21-nucleotide-long small RNAs corresponding to siRNAs. These results indicate that ALFE1-3 and AEFE1 produce piRNAs only in antisense orientation and also that these elements are located in specific regions of the genome that generate long transcripts for endogenous siRNAs and piRNA production.

**Fig 2.**
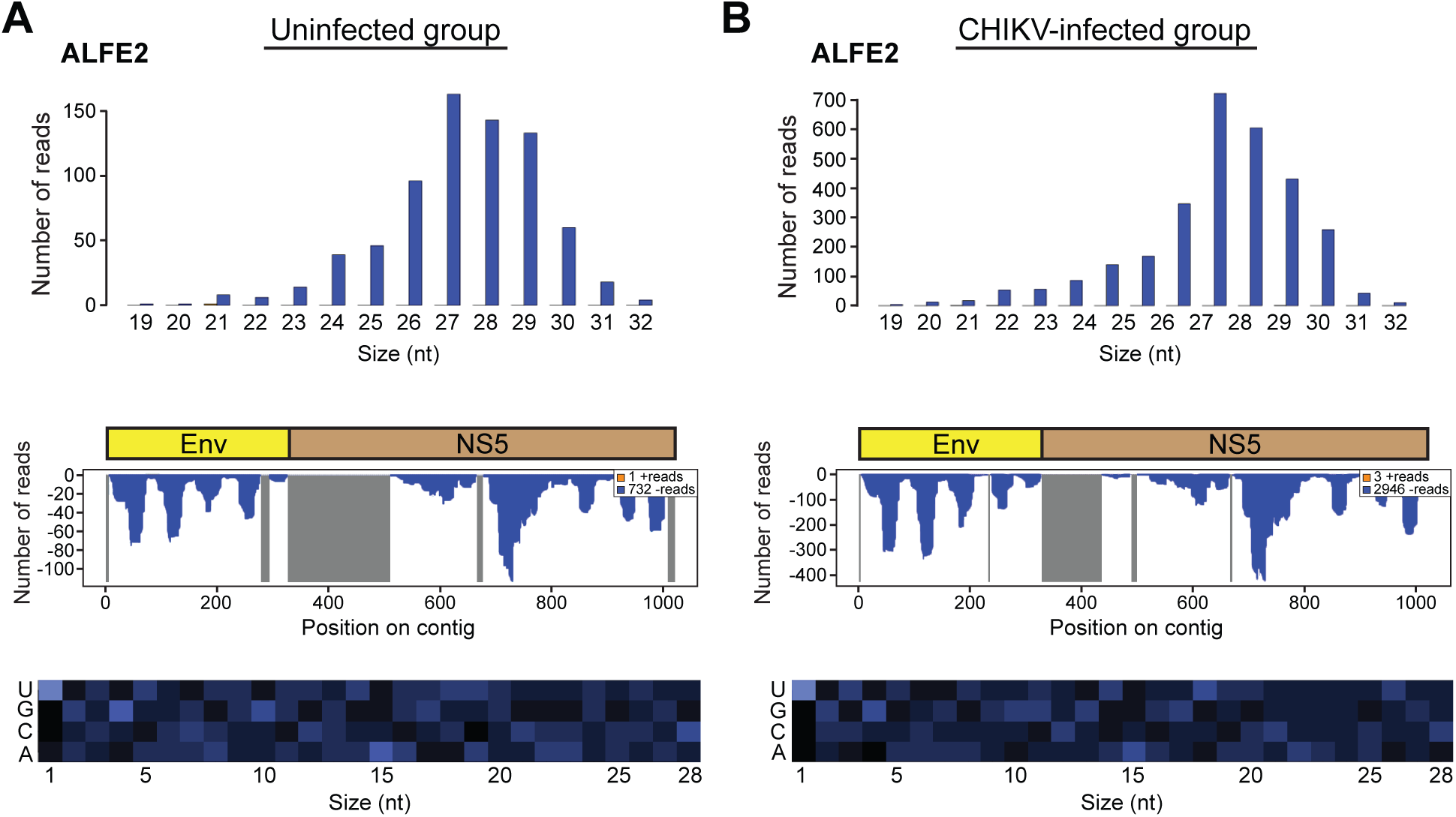
Primary piRNA-like small RNA production from ALFE2. Size distribution (upper panels) and profiles (middle panels) of small RNAs mapped to ALFE2 in *Ae. albopictus* (A) without or (B) with CHIKV infection. Orange and blue indicate positive- and negative-stranded reads, respectively. For the small RNA profiles, the x-axis represents the nucleotide position on the ALFE2-containing contig; the y-axis shows the number of reads for each nucleotide position; gray lines represent uncovered regions. Bottom panel: relative nucleotide frequency per position of the 28-nt ALFE2-derived small RNA shown as heat map. The intensity varies in correlation with frequency.

### DENV infection does not affect AEFE1 transcript abundance in *Ae. aegypti*

Virus infections affect the expression of a number of host genes, for instance DENV suppresses immune gene expression in the *Ae. aegypti* Aag2 cell line (18). The presence of mRNA and small RNAs from ALFE and AEFE in *Aedes* mosquitoes prompted us to check whether the expression level of their transcripts was altered during DENV infections *in vivo*. We utilized unpublished transcriptome datasets from *Ae. aegypti* females orally infected or uninfected with either of two DENV serotypes (DENV1 and DENV3), which were available in our laboratory to examine AEFE1 mRNA expression. DENV1 infection did not affect AEFE1 mRNA expression at 24 and 96 hours post infection (Fig. 3A). No significant difference in AEFE1 mRNA expression was observed between DENV1 and -3 infected *Ae. aegypti* at 18 and 24 hours post infection (Fig. 3B). The data suggest that AEFE1 is constitutively transcribed and not affected by DENV1 and -3 infections in *Ae. aegypti* adult females.

**Fig 3.**
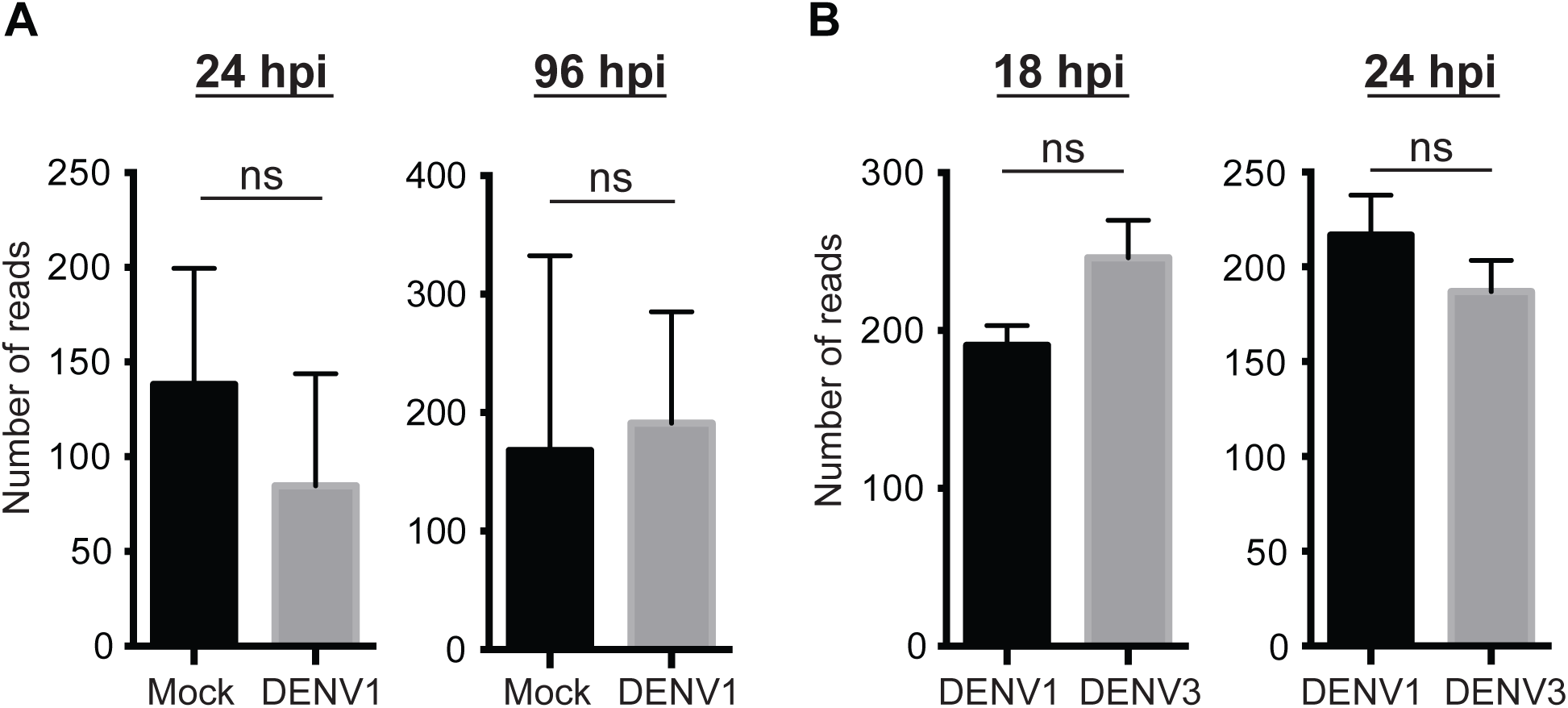
DENV1 and -3 infections have no detectable impact on AEFE1 mRNA expression *in vivo*. AEFE1 transcription level was analyzed by DESeq2 using a transcriptome dataset from *Ae. aegypti* infected (A) with or without DENV1, (B) with DENV1 or -3 at the indicated time points. The average number of reads were calculated from several individual libraries: 7 and 17 libraries for Mock and DENV1 respectively at 24 hpi; 6 and 17 libraries for Mock and DENV1 at 96 hpi; 3 for each library on DENV1 or -3 infection at both time points. Statistical significance was determined by DESeq2, ns indicates no significant difference (adjusted p-value >0.05).

### Searching for new flavivirus-like elements in *Aedes* mosquitoes

PCR for full-length ALFE1 DNA in the *Ae. albopictus* Vietnam strain did not result in amplification (Fig. 1B). We then performed PCR with primer pairs targeting ALFE1 and its flanking regions and amplified bands with unexpected sizes (data not shown). Because of the heterogeneity of the PCR products, we moved on to confirm the existence of ALFE1 and to search for new ALFEs in *Ae. albopictus*. We performed whole-genome DNA sequencing of the *Ae. albopictus* Vietnam strain and C6/36 cell line. First, we mapped reads from the *Ae. albopictus* Vietnam and C6/36 DNA libraries to ALFE1-3 and their respective flanking region sequences (Fig. 4 DNA coverage showed the presence of full-length ALFE1 in the genomes of the Vietnam strain and of C6/36 cells (Fig. 4A), despite the lack of PCR amplification. In addition, the relative coverage of ALFE1-3 flanking regions was substantially higher, indicating that these elements are integrated in multi-copy regions (Fig. 4A-C). We also observed a similar trend of insertion of ALFEs in multi-copy sequences when analyzing the genome sequence of the *Ae. albopictus* Foshan strain available in Vectorbase (data not shown).

**Fig 4.**
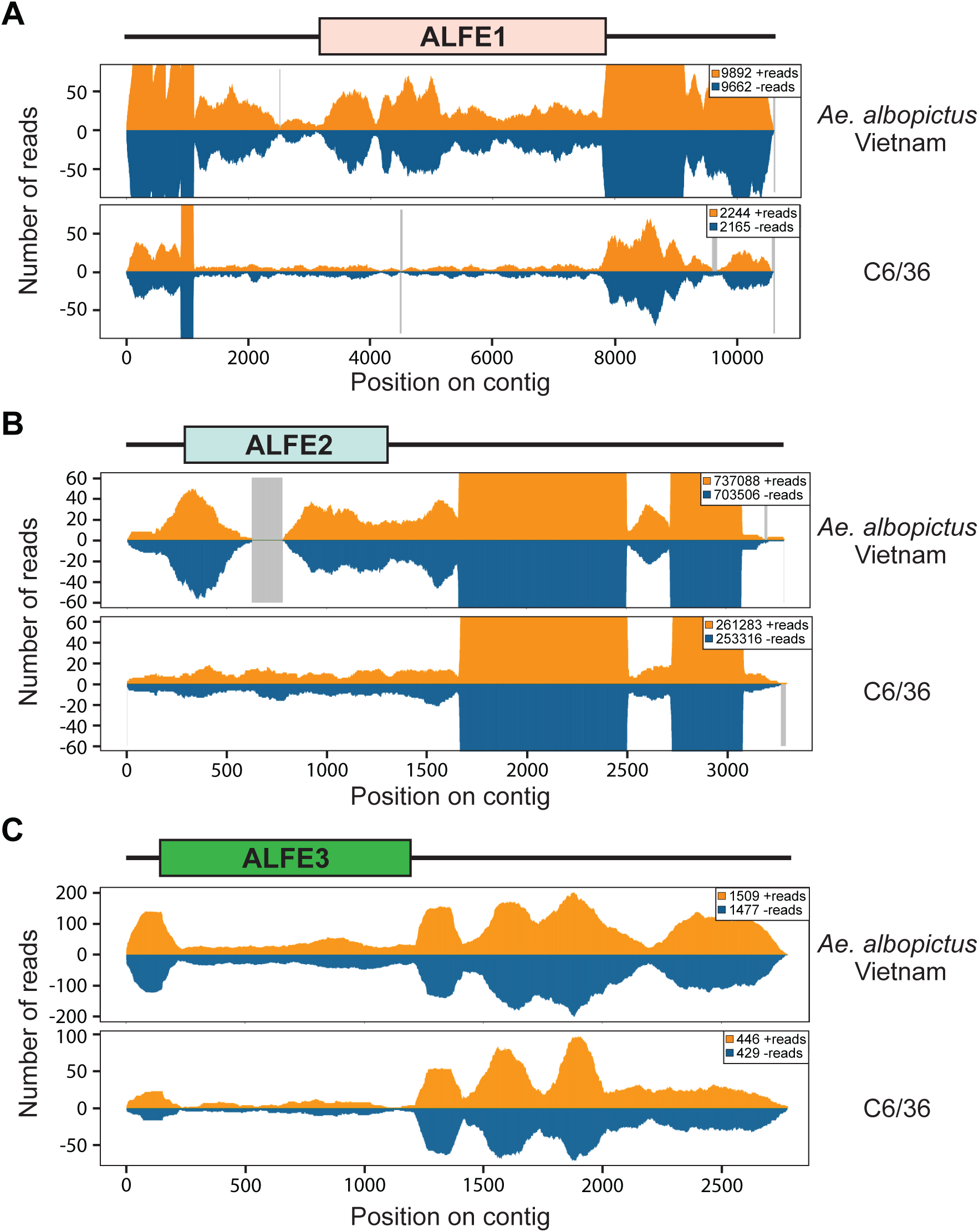
ALFE and AEFE are integrated in repetitive regions in *Aedes* mosquito genomes. DNA reads from the *Ae. albopictus* Vietnam strain and C6/36 DNA library mapped to (A) ALFE1, (B) ALFE2 and (C) ALFE3 and their flanking regions. Orange and blue indicate positive- and negative-stranded reads, respectively. Box indicates each ALFE and the black bar represents the flanking regions. The x-axis represents the nucleotide position on the ALFE-containing contig. The y-axis shows the number of reads for each nucleotide position. Gray lines represent uncovered regions.

To identify new ALFEs, we performed an *in silico* screening based on an iterative mapping and assembly procedure using ALFE1-3 sequences as scaffolds and the DNA library from the *Ae. albopictus* Vietnam strain as the query (details provided in materials and methods). This *in silico* screening yielded eight contigs named ALFE4-11 harboring ALFE1-3 partial sequences (Fig. 5A). For instance, ALFE4 was composed by a portion of ALFE1 followed by the full-length ALFE3. We detected different flanking genomic sequences to each ALFE, suggesting that multiple versions of each ALFE are present in the genome of the *Ae. albopictus* Vietnam strain. The sequences of the nearby regions are unique to *Aedes* mosquitoes, as homology search against Genbank and Vectorbase databases did not show homology with any known organisms. Moreover, the six-phase translation of the flanking sequences did not show any similarity with known proteins from UniProt Knowledge database.

**Fig 5.**
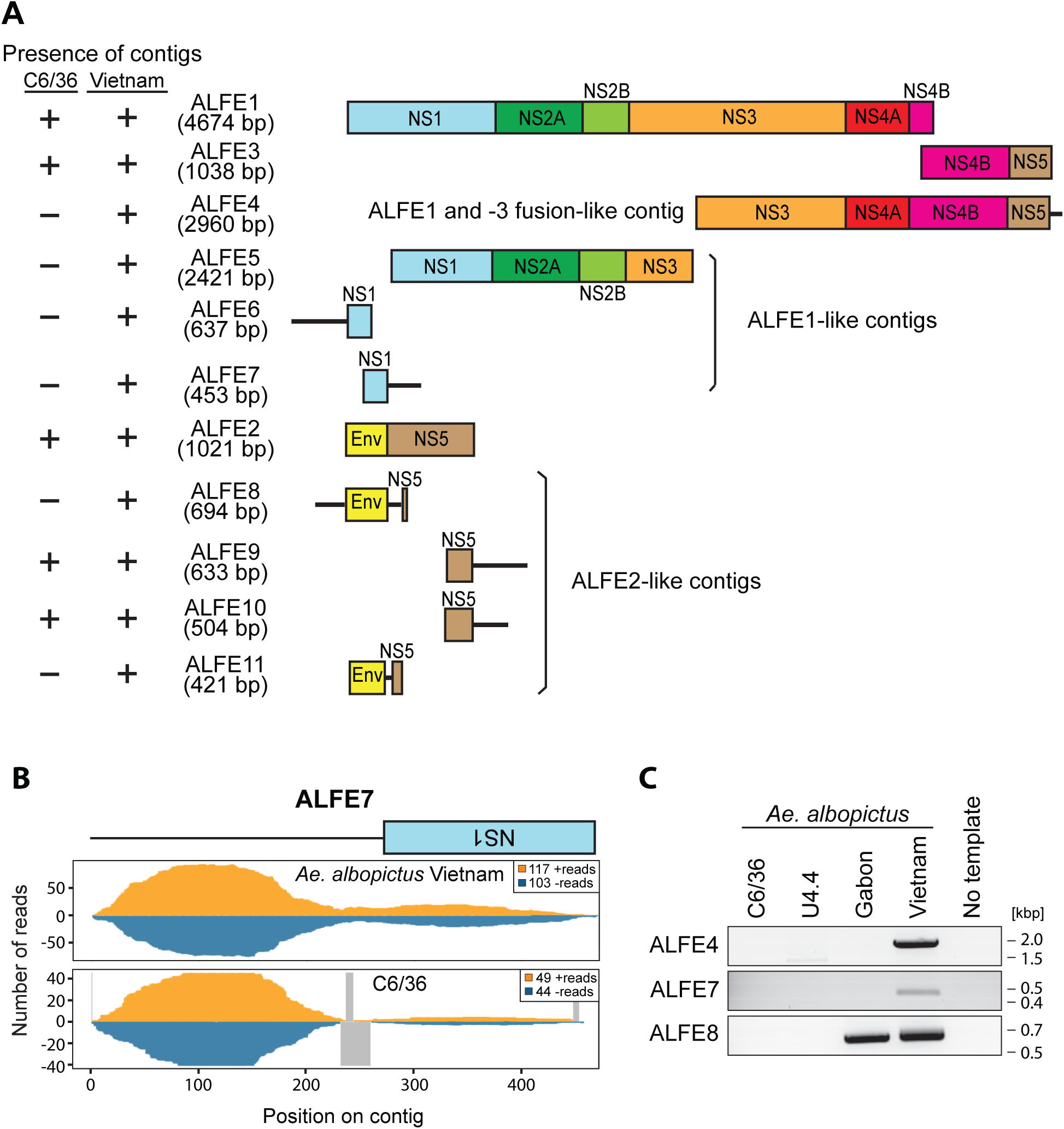
New ALFEs identified in the *Ae. albopictus* Vietnam strain. *In silico* pipeline generated 8 new ALFE-like contigs. (A) Schematic representation of ALFE-contigs. ALFE4 is a fusion element composed of ALFE1 and -3. ALFE5-7 are partial sequences of ALFE1. ALFE8-11 are ALFE2-like contigs. Box indicates ALFE-derived sequence and the black line indicates non-ALFE sequence. Presence (+) or absence (-) of the contigs is summarized on the left column. (B) DNA coverage of ALFE7 with *Ae. albopictus* Vietnam DNA library. Orange and blue indicate positive- and negative-stranded DNA respectively. Gray lines represent uncovered regions. (C) ALFE4, -7 and -8 DNA detection in C6/36 and U4.4 cell lines, and in the *Ae. albopictus* Gabon and Vietnam strains by PCR.

To confirm the existence of the bioinformatically predicted ALFE4-11, we used two different assessments: 1) quality of DNA mapping of *Ae. albopictus* Vietnam and C6/36 reads to the new ALFE contigs, and 2) PCR with primer pairs specific to the new ALFE contigs. For the DNA mapping, depth and continuity of the coverage along the ALFE sequences were used to confirm the ALFE existence in the Vietnam strain. Figure 5B shows DNA coverage and continuity of reads on ALFE7 as an example of the validation. In the *Ae. albopictus* Vietnam strain the reads continuously covered the ALFE7 contig sequence while in C6/36 cells there are differences in continuity of the coverage (presence of a gap). In addition, the read counts corresponding to the flanking regions of ALFE7 confirmed that this element is integrated in multi-copy sequences in the *Ae. albopictus* genome. Figure 5C shows PCR amplification products of predicted ALFE4, -7 and -8. These three elements were detected in the *Ae. albopictus* Vietnam strain whereas only ALFE8 was present in the Gabon strain. Neither C6/36 nor U4.4 genomes showed the presence of these elements. In addition, we checked small RNA production from the newly described ALFEs. As observed with ALFE1-3, the bioinformatically predicted ALFEs and their flanking sequences are covered by only antisense piRNA-like molecules with a U1 bias (Fig. S3).

Lastly, we compared the ALFE contigs identified from C6/36 and *Ae. albopictus* Vietnam with *Ae. albopictus* Foshan genomic DNA supercontigs. We observed considerable variation in the flanking regions between ALFEs in the different strains. For example, the *in silico* pipeline predicted the existence of ALFE4 (a fusion between ALFE1 and ALFE3) in the Vietnam strain (Fig. 6). When the ALFE4 contig was compared to ALFE4-like contig in C6/36 cells, a 1.3-kbp host sequence was present between ALFE1 and -3. This host sequence also exists in the Foshan genome but is much longer (11 kbp) and contains a repeat sequence at both extremities of a coding DNA sequence (CDS) with a gag-, reverse transcriptase- and proteinase-like domains. The analysis suggests that ALFE1 and -3 were originally the same element in the Vietnam strain (ALFE4) and were separated by insertion of a retrotransposon in the Foshan strain or vice versa.

**Fig 6.**
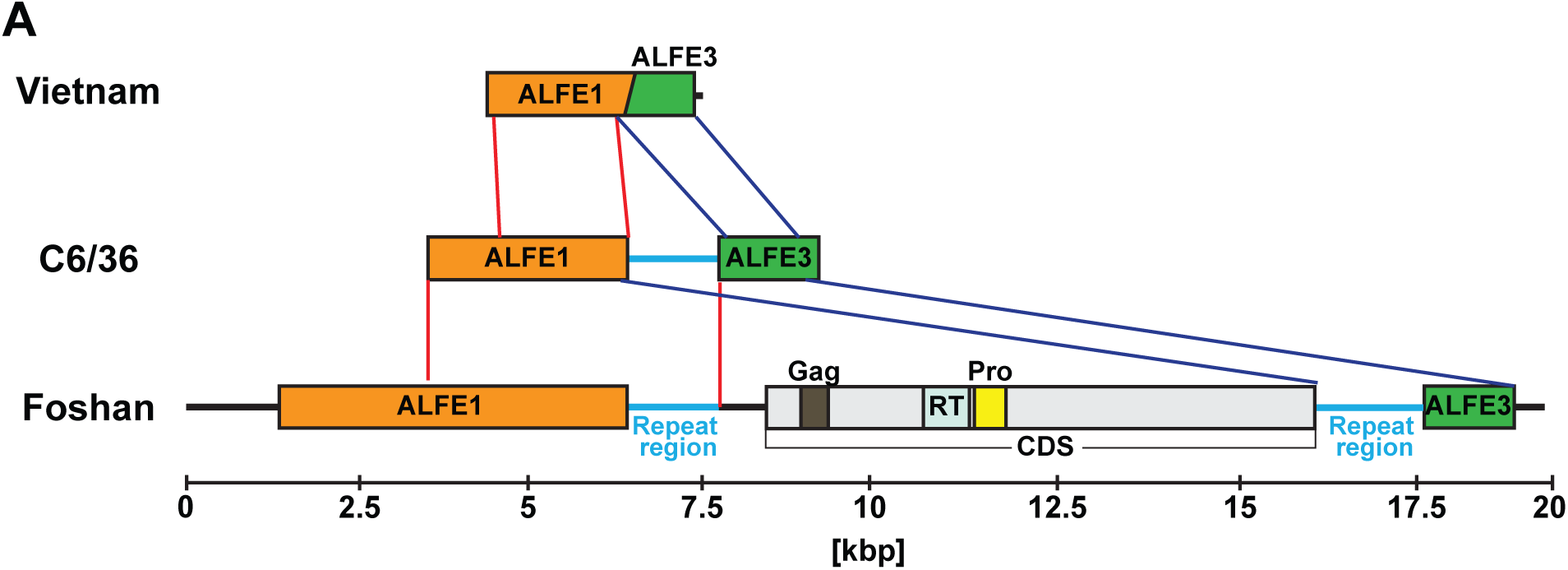
Comparison of ALFE1/3 sequence in the *Ae. albopictus* Vietnam strain, Foshan strain and C6/36 cells. Schematic representation of ALFE4 (Vietnam), ALFE4-like element (C6/36) and the Foshan ALFE1 and -3 with their flanking regions. ALFE1/3 in the Foshan strain has a gap of approximately 11 kbp in length with a CDS harboring a retrotransposon gag, reverse transcriptase and protease domains and repeat sequences at both 5’ and 3’ extremities. Overlapped regions of ALFE1 and 3 are visualized with red and blue lines. Light blue bar indicates repeat sequences.

### ALFE- and AEFE-derived proteins are undetectable by mass spectrometry

Because ALFE1-3 and AEFE1 generated mRNA *in vitro* and *in vivo*, we checked whether they could produce detectable proteins. As antibodies against ALFE, AEFE or similar flaviviruses are not available, we performed mass spectrometry (MS) analysis with C6/36 and Aag2 cell lines to find ALFE and AEFE-derived peptides. Proteins from C6/36 and Aag2 cells were purified and subjected to MS for bottom-up proteomics analysis. Although we identified Ago2, Dcr2 and Piwi5, as well as thousands of proteins from all the subcellular fractions and ranging from 5.6 to 811 kDa (Table 1 and Table S1), no ALFE- neither AEFE-proteolytic peptides were identified in both C6/36 and Aag2 cells. This result suggests that ALFE1-3 and AEFE1 are not translated at high level, if translate at all, in both mosquito cell lines.

**Table 1.** Proteomic characterization of C6/36 and Aag2 cell lines.

|  | EFVEs | Piwi5 | Dcr2 | Ago2 | Number of identified proteins |
| --- | --- | --- | --- | --- | --- |
| C6/36 | 0 | 3 | 0* | 2 | 1942 ± 637 |
| Aag2 | 0 | 3 | 2 | 2 | 2054 ± 291 |

Number of times that proteolytic peptides corresponding to EVEs proteins and positive controls (out of 3 experiments) were found. The average total number of identified proteins is depicted. \*C6/36 cells are Dcr2 deficient.

### Phylogenetic analysis of AEFE1 in *Ae. aegypti*

As mentioned above, different environmental conditions or reduced gene flow could result in divergent evolution of EVEs between geographical locations. To study this, we examined AEFE1 in *Ae. aegypti* from Cameroon, French Guiana and Thailand. The full-length sequence of AEFE1 in individual mosquitoes from each strain was amplified and sequenced. Some individual mosquitoes from Cameroon and Thailand populations did not show amplification of AEFE1 and were therefore excluded from this analysis. Next, we generated maximum-likelihood phylogenetic trees of AEFE1 consensus sequences from each individual and their homologous sequence in the genome of ISFs closely related to AEFE1 such as CFAV, KRV and AEFV, as well as homologous sequences in related medically important flaviviruses. As expected, the AEFE1 sequence is closely related to the ISFs (Fig. S4). To further examine the relationship among AEFE1 sequences from different geographical locations, an additional tree was generated and rooted with the ISFs as the outgroup, but is presented without the ISFs for visual clarity (Fig. 7). The phylogenetic analysis showed that among the three ISFs considered, KRV is the closest to AEFE1 (Fig. S4). In addition, the phylogenetic relationships among AEFE1 sequences mainly recapitulate the recent evolutionary history of *Ae. aegypti* populations (Fig. 7). With two exceptions, AEFE1 sequences in mosquitoes from the non-African (Thailand and French Guiana) populations were derived from AEFE1 sequences found in the African (Cameroon) population. A basal evolutionary position of African populations is typically observed for *Ae. aegypti* genes (19), consistent with the recent out-of-Africa geographical expansion of the species. Thus, the phylogenetic analysis suggests that AEFE1 evolved consistently with other host genes at the species level.

**Fig 7.**
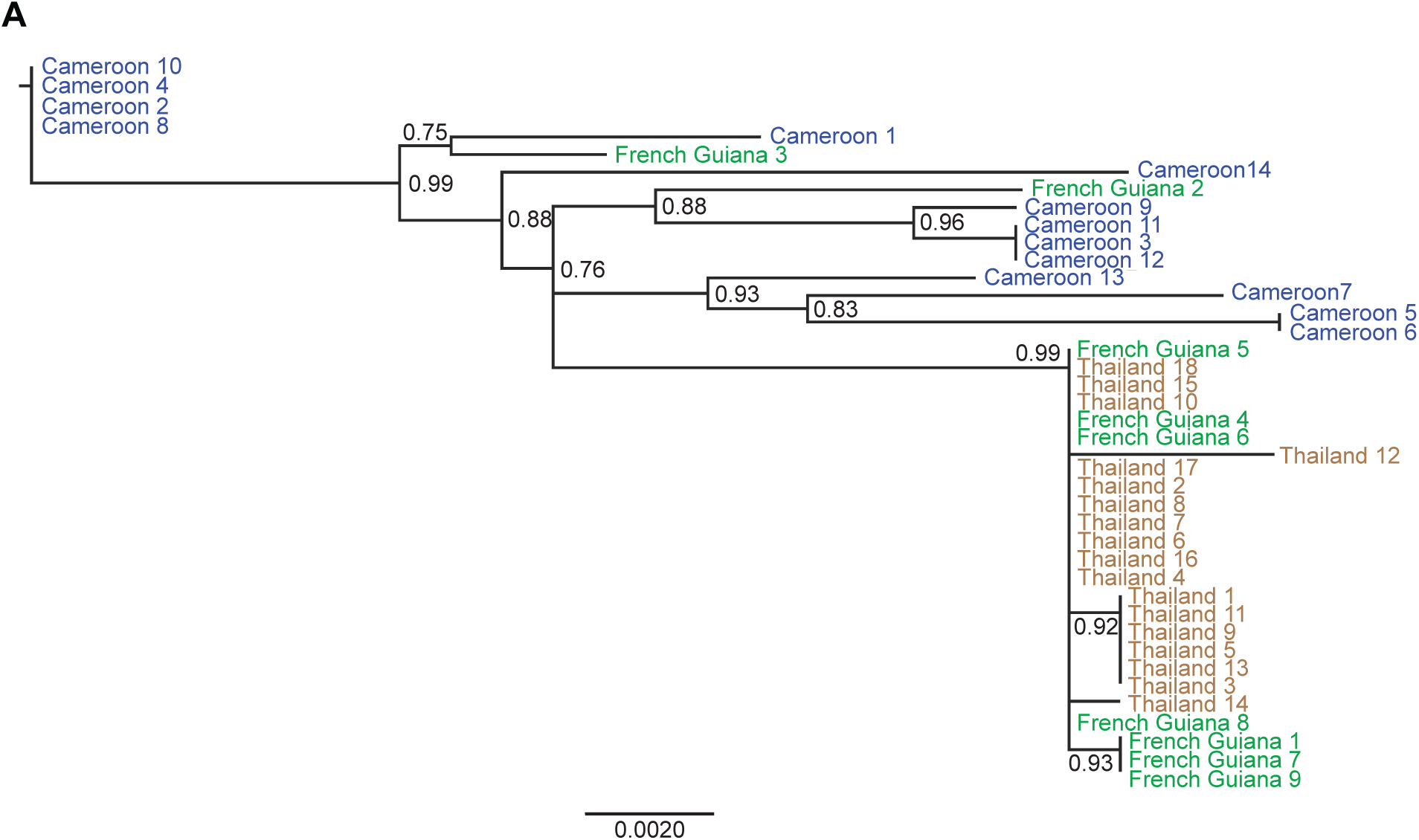
Phylogenetic relationships among AEFE1 sequences from different *Ae. aegypti* populations. A maximum-likelihood tree was generated using the Fast likelihood-based approach. AEFE1 DNA was sequenced in individual mosquitoes from the *Ae. aegypti* Cameroon, French Guiana and Thailand strains. The scale bar indicates the number of nucleotide substitutions per site and node support values are shown for major nodes. The tree was rooted with ISF sequences (S4 Fig) but the outgroup was omitted for visual clarity. Node values represent Shimodaira-Hasegawa (SH)-like branch support (only values > 0.7 are shown).

## Discussion

Recently, reactivation of EVEs due to related or unrelated viral infections have been reported, strongly suggesting a role of EVEs during the immune response of the host (9-11). Dozens of EVEs, comprising flavivirus-, rhabdovirus- and reovirus-related EVEs have been bioinformatically predicted for different mosquito strains and some of them were confirmed in mosquito cell lines or strains (4, 5, 7, 8, 20). However, few studies have been conducted to characterize EVEs *in vivo*, a necessary step to further investigate their role during viral infection of mosquitoes. Due to the current disease outbreaks caused by mosquito-borne flaviviruses, such as Zika and dengue viruses, we decided to study and characterize *in vivo* endogenous flaviviral elements (EFVEs) that were previously identified in *Ae. albopictus* and *Ae. aegypti* mosquito cell lines. To improve our understanding of the forces shaping the evolution of these EFVEs, we assessed their presence, transcription, small RNA production, protein production and phylogeny using African, Asian and American populations of *Aedes* mosquitoes. These wild-type populations are epidemiologically relevant because they occur in regions where they act as the main arbovirus vectors (21, 22). As to simplify nomenclature, we propose to name ALFEx the EFVEs identified in *Ae. albopictus* and AEFEx the ones identified in *Ae. aegypti*, where x is a number.

We first showed the presence of full-length ALFE1-3 and AEFE1 DNA in *Ae. albopictus* or *Ae. aegypti* individuals, respectively, in several populations of both mosquito species. This indicates that ALFE1-3 and AEFE1 are likely derived from ancient integration events of flavivirus DNA in nature that have persisted during recent evolution of the species. We also detected complete or partial mRNA of ALFE1-3 and AEFE1 *in vivo*. The expression level of AEFE1 remained unchanged following DENV1 and -3 infection of *Ae. aegypti* mosquitoes, suggesting a constitutive expression. However, experiments addressing AEFE and ALFE mRNA regulation during different virus infections or different abiotic or biotic stimuli should be designed and performed in the future. In most documented cases, functional EVEs play a role at the protein level (9-12). We performed a powerful bottom-up proteomic approach to search for ALFE- and AEFE- proteolytic peptides in C6/36 and Aag2 cell lines. The mass spectrometry analysis could not validate the expression of ALFE1-3 and AEFE1 proteins in any of the subcellular fractions analyzed, while control peptides for each cell type were readily detected. Due to the conservation of ALFE and AEFE *in vivo* and *in vitro*, this result strongly suggests that ALFE and AEFE mRNAs are not translated into proteins. However, EFVEs could need specific conditions to be translated. For instance, the mosquito cell lines we used were established from larvae or embryos (23, 24). ALFE and AEFE could be translated only in specific tissues or stages of development *in vivo*. Another interesting possibility is that because the transcripts were observed by RT-PCR and RNA sequencing, ALFE and AEFE could have a function at the RNA levels.

Small RNAs have critical roles in various aspects of host fitness and antiviral immunity in mosquitoes (16, 25-31). The piRNA pathway is one of the small RNA pathways and is mainly controlling “non-self” sequences, such as retrotransposable elements (32-34). According to the PIWI proteins involved and available for piRNA biogenesis, they are classified into primary piRNAs or secondary piRNAs, the latter being produced from an amplification cycle known as the ping-pong cycle. We found that all ALFE1-3, AEFE1 and their flanking regions produced a population of primary piRNA-like molecules only in antisense orientation that were not affected during CHIKV infection. A recent study also found that some annotated genes produce antisense primary piRNA-like small RNAs in *Ae. aegypti* Aag2 cells (35). Two hypotheses could account for the production of only primary piRNA-like molecules by ALFE and AEFE. First, the production of ALFE- and AEFE-derived primary piRNAs may be only happening in specific tissues lacking PIWI proteins, such as Piwi5 and/or -6, involved in ping-pong amplification. Second, there could exist an unknown selection mechanism to produce only primary piRNA from EVE- enriched regions. Even if these hypotheses could account for the only primary piRNAs, none of them explains why piRNAs are only in antisense orientation. Curiously, in vertebrates, some murine EBLNs and their flanking regions also generate antisense piRNA-like small RNAs (36). This suggest that EVEs have common features on small RNA productions across species and kingdoms. To elucidate how and why this is happening deserver further studies.

Regarding the function of these EFVE-derived piRNAs, it is tempting to propose that they could regulate infections by closely related viruses by targeting viral RNA in *Aedes* mosquitoes. ALFE and AEFE do not contain sequences of 24-30 nucleotides that perfectly span AEFV and KRV sequences, which are the most closely related known ISFs. However, piRNAs allow some mismatches with target sequences (37-39). If ALFE and AEFE-derived piRNAs are loaded into the ping-pong amplification cycle during viral infection, a number of piRNAs that match the virus could be produced and those piRNAs could contribute to control viral replication. We and others previously demonstrated that viral piRNAs (vpiRNAs) were detected in *Aedes* mosquitoes and in mosquito cell lines infected with mosquito-borne viruses such as CHIKV and Rift Valley Fever virus (phlebovirus, *Bunyaviridae* family) (16, 28). Although the direct effects of vpiRNA on viral replication remains unclear, vpiRNAs could render the mosquitoes tolerant to the virus infections (16). By providing closely related piRNAs into the ping-pong cycle, ALFE and AEFE may contribute to control pathogenesis during exogenous viral infections.

Using a bioinformatics approach, we identified several new ALFEs composed of partial or complete ALFE1-3 sequences in *Ae. albopictus* mosquitoes from Vietnam. Some of these new ALFEs were predicted only in our mosquito strain but not found in the C6/36 cell line genome. PCR also showed variability in the detection of ALFE4-11 among the mosquito strains, suggesting that each mosquito population has a different set of EFVEs in their genome. The recent genome sequencing of *Ae. albopictus* Foshan strain *Chen et al*. (8) found more than 20 EFVEs that were mostly related to NS1 or NS5 genes from ISFs. Likewise, we also found three elements containing NS1 and four containing NS5 out of eight identified ALFE-like contigs. Another study performed on field-collected *Ae. albopictus* in Northern Italy detected flaviviral NS5 related to CFAV and KRV NS5 (40). Therefore, *Aedes* mosquitoes seem to have preferentially accumulated ancient flaviviral NS5 sequences in their genome. There are two potential and not exclusive explanations for this: (1) NS5 could be preferably endogenized compared to other flaviviral sequences, (2) NS5 sequences could be positively maintained over generations. Although the mechanism of endogenization has not been elucidated, we and others suggested that it happens in a retrotransposon-dependent manner (16, 41, 42). *Ae. albopictus* and *Ae. aegypti* genomes harbor 50% to 70% of repetitive sequences such as retrotransposons and long interspersed nuclear elements (LINEs) (8, 43). Accordingly, it is interesting to propose that the sequence of NS5 could be efficiently recognized by retrotransposons in *Aedes* mosquito genomes.

One interesting observation arose when we compared the newly identified ALFE4 between our Vietnam strain and the Foshan strain of *Ae. albopictus*. In the Vietnam strain, ALFE4 appears as a fusion of ALFE1 and ALFE3. In the Foshan strain, a large CDS containing retrotransposon RT-like domain with terminal repeat sequences exists between ALFE1 and -3. Katzourakis *et al*. (5) observed in *Ae. aegypti* and *Ae. albopictus* the presence of an almost entire flaviviral genome fragmented in several pieces across the mosquito genome. Likely, we propose that ALFE- and AEFE-like sequences are inserted in repetitive regions of the mosquito genome, where transposons can be acting as a trap for non-retroviral DNA. These intergenic flanking regions display substantial structural variation among mosquito populations, arguing for an active rearrangement and continuous change of EFVE-containing regions and pointing to a role of these captured viral sequences in the evolution of host genomes.

It is, however, important to stress that we faced significant challenges working with the genome sequence of the *Ae. albopictus* Vietnam strain. This is presumably due to the difficulties in assembling the different contigs, despite a comfortable depth of coverage, due to the high content of repetitive sequences. Indeed, the currently available genome assemblies of *Ae. albopictus* and *Ae. aegypti* in Vectorbase consist of thousands of unassembled supercontigs. There is an urgent need to assemble these fragmented reference genome sequences into end-to-end chromosome maps. Hopefully, the advent of new tools such as long-read sequencing technologies and chromosome confirmation capture will be a leap forward in the study of mosquito genomes.

Finally, a phylogenetic analysis of AEFE1 in *Ae. aegypti* from Cameroon, French Guiana and Thailand showed that the evolutionary history of AEFE1 was similar to that of most nuclear genes (19). AEFE1 was genetically more diverse in the Cameroon strain compared to the French Guiana and Thailand strains, which is also consistent with patterns of genetic diversity for nuclear genes. Our phylogenetic analysis suggests that the integration events of AEFE1 occurred before *Ae. aegypti* mosquitoes expanded out of the African continent. However, we found some *Ae. aegypti* individuals from Cameroon and Thailand in which the full-length AEFE1 could not be amplified, suggesting that both structural and nucleotide variants of AEFE1 exist within the same population. It is likely that following the initial integration event, complex evolutionary forces have shaped the genetic diversity of EFVEs that is observed today. Unraveling this diversity will be necessary to elucidate their potential functional role.

## Materials and Methods

### Ethics statement

The Institut Pasteur animal facility has received accreditation from the French Ministry of Agriculture to perform experiments on live animals in compliance with the French and European regulations on care and protection of laboratory animals. Rabbit blood draws performed during the course of this study were approved by the Institutional Animal Care and Use Committee at Institut Pasteur under protocol number 2015-0032, in accordance with European directive 2010/63/UE and French legislation.

### Cell culture

C6/36 (ATCC CRL-1660) and U4.4 (Ae. *albopictus*) (24), and Aag2 (Ae. *aegypti*) (44) cell lines (kindly provided by G. P. Pijlman, Wageningen University, the Netherlands) were maintained 28°C in L-15 Leibovitz medium (Gibco) supplemented with 10% foetal bovine serum (Gibco), 1% nonessential amino acids (Gibco), 2% tryptose phosphate broth (Sigma) and 1% penicillin/streptomycin (Gibco).

### Mosquitoes

Laboratory colonies of *Ae. aegypti* were established from field collections in Cameroon (2014), French Guiana (2015), and Thailand (2013). Laboratory colonies of *Ae. albopictus* were established from field collections in Gabon (2014) and Vietnam (2011). All the experiments were performed within 16 generations of laboratory colonization. The insectary conditions for mosquito maintenance were 28°C, 70% relative humidity and a 12 h light: 12 h dark cycle. Adults were maintained with permanent access to 10% sucrose solution. Adult females were offered commercial rabbit blood (BCL) twice a week through a membrane feeding system (Hemotek Ltd).

### Experimental DENV infections *in vivo*

Wild-type, low-passage DENV isolates (DENV1: KDH0030A (45); DENV3: GabonMDA2010 (46)) were originally obtained in 2010 from the serum of dengue patients. Informed consent of the patients was not necessary because the viruses isolated in cell culture are longer considered human samples. KDH0030A was isolated at the Armed Forces Research Institute of Medical Sciences, Bangkok, Thailand. GabonMDA2010 was isolated at the Centre International de Recherches Médicales de Franceville, Gabon. Virus stocks were prepared and experimental mosquito infections were conducted as previously described (47). Briefly, 4- to 7-day-old female mosquitoes were deprived of sucrose for 24 hours and transferred to a biosafety level-3 insectary. Washed rabbit erythrocytes resuspended in PBS were mixed 2:1 with pre-diluted viral stock and supplemented with 10 mM ATP (Sigma-Aldrich). The viral stock was prediluted in Leibovitz’s L-15 medium with 10% FBS, 0.1% penicillin/streptomycin and 1% sodium bicarbonate (Gibco) to reach an infectious titer ranging from 5 x 10^6^ to 1.1 x 10^7^ focus-forming units per mL of blood using a standard focus-forming assay in C6/36 cells (47). A control blood meal was prepared identically except that the supernatant of mock-inoculated cells replaced the viral suspension. Mosquitoes were offered the infectious or control blood meal for 30 min through a membrane feeding system (Hemotek Ltd) set at 37°C with a piece of desalted pig intestine as the membrane. Following the blood meal, fully engorged females were selected and incubated at 28°C, 70% relative humidity and under a 12 h light: 12 h dark cycle with permanent access to 10% sucrose.

### PCR and RT-PCR

Total DNA was extracted from mosquito cell lines or individuals with NucleoSpin Tissue kit (Macherey-Nagel). Total RNA was extracted from the mosquito samples with TRIzol (Invitrogen). Following DNase I (Promega) treatment, cDNA was synthesized with Maxima H Minus Reverse Transcriptase and oligodT primers according to the manufacturer’s instructions (Thermo Fisher Scientific). PCR and nested PCR were performed with DreamTaq DNA polymerase (Thermo Fisher Scientific). The sequences of the primers are provided in supplementary Table S2.

### Transcriptome data analysis

Individual midgut libraries of *Ae. aegypti* infected with DENV-1 or DENV-3 were prepared from total RNA extracts from individual midguts after quality control with a Bioanalyzer RNA 6000 kit (Agilent). Purification and fragmentation of mRNA, cDNA synthesis, end-repair, A-tailing, Illumina indexes ligation and PCR amplification were performed using TruSeq RNA Sample Prep v2 (Illumina) followed by cDNA quality check by Bioanalyzer DNA 1000 kit (Agilent). Libraries were diluted to 10 pM after Qubit quantification (Thermo Fisher Scientific), loaded onto a flow cell, clustered with cBOT (Illumina). Single-end reads of 51 nucleotides in length were generated on a HiSeq2000 sequencing platform (Illumina). Sequencing reads with a quality score <30 were trimmed using Cutadapt (https://cutadapt.readthedocs.io/en/stable/). Passing-filter reads were mapped to *Ae. aegypti* transcripts (AaegL3.1,http://vectorbase.org) using Bowtie2 then processed with the Samtools suite to create of a matrix of raw counts used for gene expression analysis by DESeq2 package.

### Small RNA and genome DNA libraries

To analyze small RNA productions from ALFE and AEFE, we used small RNA libraries of *Ae. albopictus* and *Ae. aegypti* infected with or without CHIKV that are publically available in the Sequence Read Archive with accession code SRP062828. For genomic DNA libraries, total DNA from C6/36 and *Ae. albopictus* Vietnam was extracted using NucleoSpin Tissue kit (Macherey-Nagel). Genomic DNA was then sheared into 200 bp using Covaris S220 device with the following parameters: Peak Incident Power, 175; Duty Factor, 10; Cycle Burst, 200; Duration, 180 seconds. Genomic DNA libraries were prepared using KAPA LTP Libray Preparation Kit Illumina Platforms (KAPA BIOSYSTEMS). The library was amplified with 10 PCR cycles and 2 x 151 pair-end reads were sequenced on NextSeq 500.

For bioinformatics analysis, the quality of fastq files was assessed with FastQC (www.bioinformatics.babraham.ac.uk/projects/fastqc/). Low-quality bases and adaptors were trimmed from each read by Cutadapt. Only reads with acceptable quality (phred score 20) were retained. A second set of graphics was generated by FastQC using the fastq files trimmed by cutadapt. Reads were mapped to target sequences using bowtie1 with the ‘-v 1’ or bowtie2 with the ‘‐‐sensitive’ for small RNA or DNA library, respectively. Bowtie1 (small RNA library) and -2 (DNA library) generate results in sam format. All sam files were analyzed by the samtools package to produce bam indexed files. Home-made R scripts with Rsamtools and Shortreads in Bioconductor were used for analysis of the bam files.

### *In silico* screening for new EFVEs

To identify all versions of EVEs in the mosquito DNA contigs we developed an iterative bioinformatics pipeline. Each iteration is composed of 4 steps:

1/ Blastn using DNA library as database and with an e-value threshold of 1E-20
2/ Reads extraction according to the blastn result
3/ Assembly of these reads using SPAdes 3
4/ Extraction of contigs larger than 400 bases

For the first iteration, the already known ALFEs sequences were used as queries. For the following iterations, the contigs selected from previous iteration were used as queries. Due to the very large number of matches detected by blastn after some iterations (repetitive regions), only 5 iteration were used in order to analyze the contigs obtained.

### Mass Spectrometry

C6/36 and Aag2 cells (10^7^) were lysed by sonication (twice 20 sec using an ultrasonic probe) in lysis buffer (urea 6M; TrisHCl 150 mM pH 8.8; β octyl 1%; DTT 10 mM). In addition, in order to have an indication of subcellular location of the proteins of interest, subcellular fractionation of cells extract was performed using a commercial kit (Subcellular Protein Fractionation Kit for Cultured Cells, Thermo Fisher Scientific), according to manufacturer’s instructions, resulting in 6 fractions: cytoplasmic, membrane, soluble nuclear, chromatin-bound, cytoskeletal extracts, and insoluble pellet. Proteolysis was performed using Filter-Assisted Sample Preparation strategy. Briefly, proteins were transferred over a 10 kDa filter (Microcon, Amicon Merck), reduced and alkylated (DTT 10 mM final, 2h, 37 °C; iodoacetamide, 50 mM final, 30 min in dark at room temperature). After 3 times-washing with ammonium bicarbonate 50 mM, proteins were proteolysed with trypsin (10 ng modified sequencing grade trypsin; Roche; 37°C, overnight). The resulting proteolytic peptides were recovered by centrifugation (15 min at 10,000 x g), acidified with 0.1% aqueous TFA and desalted using C18 Sample Prep Pipette Tips (Ziptip C18, Millipore). Peptides were purified on a capillary reversed-phase column (C18 Acclaim PepMap100 Å, 75 μm i.d., 50-cm length; Thermo Fisher Scientific) at a constant flow rate of 220 nL/min, with a gradient of 2% to 40% buffer B in buffer A in 170 min; buffer A: H_2_O/acetonitrile(ACN)/ FA 98:2:0.1 (v/v/v); buffer B: H_2_O/ACN/FA 10:90:0.1 (v/v/v). The MS analysis was performed on a Q Exactive mass spectrometer (Thermo Fisher Scientific) with a top 10 acquisition method: MS resolution 70,000, mass range 400–2,000 Da, followed by 10 MS/MS on the ten most intense peaks at resolution 17,500, with a dynamic exclusion for 90 s. Raw data was processed using Proteome Discoverer 2.1 (Thermo Fisher Scientific). The database search was done with Mascot search engine (Matrix Science Mascot 2.2.04) on a home-made protein databank containing the putative proteins for endogenous viral elements as well as *Aedes* proteins (17,756 sequences). The following parameters were used: MS tolerance 10 ppm; MS/MS tolerance 0.02 Da; semi-tryptic peptides; two miscleavages allowed; partial modifications carbamidomethylation ©, oxidation (M), deamidation (NQ).

### Phylogenetic analysis of AEFE1

DNA was extracted from individual *Ae. aegypti*. PCR was performed for the full-length AEFE1 element and the amplicons were sequenced by the Sanger technique. Forward and reverse sequences were trimmed based on chromatogram quality and aligned to generate a consensus sequence using the program Geneious v7 (http://www.geneious.com, (48)). Sequences from all successfully sequenced individuals, closely related ISFs, and medically important flaviviruses were aligned using ClustalW and trimmed to the same length. The program PHYML (49) was used to generate two phylogenetic trees using the PhyML Best AIC Tree and the Fast likelihood-based method. The first tree contains closely related ISFs, medically relevant flavivirus, and a representative AEFE1 sequence. The second tree was constructed using only the ISFs and the AEFE1 sequences from *Ae. aegypti*. The best nucleotide substitution method was GTR +G for both trees.

## Acknowledgements

We thank all members of the Saleh and Lambrechts labs for fruitful discussions; Catherine Lallemand for assistance with mosquito rearing; and Valerie Dorey for help with preparation of DNA libraries. We are grateful to Christophe Paupy, Diego Ayala, Davy Jiolle, Elisée Nchoutpouen, Karima Zouache, Alongkot Ponlawat, Thanyalak Fansiri and Isabelle Dusfour for the original mosquito colonies.

This work was supported by the European Research Council (FP7/2013-2019 ERC CoG 615220 to MCS), the French Government’s Investissement d’Avenir program, Laboratoire d’Excellence Integrative Biology of Emerging Infectious Diseases (grant ANR-10-LABX-62-IBEID to LL and MCS), the City of Paris Emergence(s) program in Biomedical Research (to LL), the European Union’s Horizon 2020 research and innovation programme under ZikaPLAN grant agreement No 734584 (to LL), and the Conseil Régional Ile de France (Sesame2010 to JV). YS is supported by Roux-Cantarini Postdoctoral Fellowships and Japan Society for the Promotion of Science Postdoctoral Fellowships for Research Abroad. The funders had no role in study design, data collection and analysis, decision to publish, or preparation of the manuscript.

## References

1. Holmes EC. 2011. The evolution of endogenous viral elements. Cell Host Microbe 10:368–377.

2. Bejarano ER, Khashoggi A, Witty M, Lichtenstein C. 1996. Integration of multiple repeats of geminiviral DNA into the nuclear genome of tobacco during evolution. Proc Natl Acad Sci U S A 93:759–764.

3. Horie M, Honda T, Suzuki Y, Kobayashi Y, Daito T, Oshida T, Ikuta K, Jern P, Gojobori T, Coffin JM, Tomonaga K. 2010. Endogenous non-retroviral RNA virus elements in mammalian genomes. Nature 463:84–87.

4. Crochu S, Cook S, Attoui H, Charrel RN, De Chesse R, Belhouchet M, Lemasson JJ, de Micco P, de Lamballerie X. 2004. Sequences of flavivirus-related RNA viruses persist in DNA form integrated in the genome of Aedes spp. mosquitoes. J Gen Virol 85:1971–1980.

5. Katzourakis A, Gifford RJ. 2010. Endogenous viral elements in animal genomes. PLoS Genet 6:e1001191.

6. Belyi VA, Levine AJ, Skalka AM. 2010. Unexpected inheritance: multiple integrations of ancient bornavirus and ebolavirus/marburgvirus sequences in vertebrate genomes. PLoS Pathog 6:e1001030.

7. Fort P, Albertini A, Van-Hua A, Berthomieu A, Roche S, Delsuc F, Pasteur N, Capy P, Gaudin Y, Weill M. 2012. Fossil rhabdoviral sequences integrated into arthropod genomes: ontogeny, evolution, and potential functionality. Mol Biol Evol 29:381–390.

8. Chen XG, Jiang X, Gu J, Xu M, Wu Y, Deng Y, Zhang C, Bonizzoni M, Dermauw W, Vontas J, Armbruster P, Huang X, Yang Y, Zhang H, He W, Peng H, Liu Y, Wu K, Chen J, Lirakis M, Topalis P, Van Leeuwen T, Hall AB, Jiang X, Thorpe C, Mueller RL, Sun C, Waterhouse RM, Yan G, Tu ZJ, Fang X, James AA. 2015. Genome sequence of the Asian Tiger mosquito, Aedes albopictus, reveals insights into its biology, genetics, and evolution. Proc Natl Acad Sci U S A 112:E5907–5915.

9. Fischer MG, Hackl T. 2016. Host genome integration and giant virus-induced reactivation of the virophage mavirus. Nature 540:288–291.

10. Fujino K, Horie M, Honda T, Merriman DK, Tomonaga K. 2014. Inhibition of Borna disease virus replication by an endogenous bornavirus-like element in the ground squirrel genome. Proc Natl Acad Sci U S A 111:13175–13180.

11. Best S, Le Tissier P, Towers G, Stoye JP. 1996. Positional cloning of the mouse retrovirus restriction gene Fv1. Nature 382:826–829.

12. Anderson MM, Lauring AS, Burns CC, Overbaugh J. 2000. Identification of a cellular cofactor required for infection by feline leukemia virus. Science 287:1828–1830.

13. Cammisa-Parks H, Cisar LA, Kane A, Stollar V. 1992. The complete nucleotide sequence of cell fusing agent (CFA): homology between the nonstructural proteins encoded by CFA and the nonstructural proteins encoded by arthropod-borne flaviviruses. Virology 189:511–524.

14. Hoshino K, Isawa H, Tsuda Y, Sawabe K, Kobayashi M. 2009. Isolation and characterization of a new insect flavivirus from Aedes albopictus and Aedes flavopictus mosquitoes in Japan. Virology 391:119–129.

15. Crabtree MB, Sang RC, Stollar V, Dunster LM, Miller BR. 2003. Genetic and phenotypic characterization of the newly described insect flavivirus, Kamiti River virus. Arch Virol 148:1095–1118.

16. Goic B, Stapleford KA, Frangeul L, Doucet AJ, Gausson V, Blanc H, Schemmel-Jofre N, Cristofari G, Lambrechts L, Vignuzzi M, Saleh MC. 2016. Virus-derived DNA drives mosquito vector tolerance to arboviral infection. Nat Commun 7:12410.

17. Burt FJ, Chen W, Miner JJ, Lenschow DJ, Merits A, Schnettler E, Kohl A, Rudd PA, Taylor A, Herrero LJ, Zaid A, Ng LF, Mahalingam S. 2017. Chikungunya virus: an update on the biology and pathogenesis of this emerging pathogen. Lancet Infect Dis doi:10.1016/s1473-3099(16)30385-1.

18. Sim S, Dimopoulos G. 2010. Dengue virus inhibits immune responses in Aedes aegypti cells. PLoS One 5:e10678.

19. Brown JE, Evans BR, Zheng W, Obas V, Barrera-Martinez L, Egizi A, Zhao H, Caccone A, Powell JR. 2014. Human impacts have shaped historical and recent evolution in Aedes aegypti, the dengue and yellow fever mosquito. Evolution 68:514–525.

20. Lequime S, Lambrechts L. 2017. Discovery of flavivirus-derived endogenous viral elements in Anopheles mosquito genomes supports the existence of Anopheles-associated insect-specific flaviviruses. Virus Evol 3:vew035.

21. Forum on Microbial T, Board on Global H, Health, Medicine D, National Academies of Sciences E, Medicine. 2016. The National Academies Collection: Reports funded by National Institutes of Health, Global Health Impacts of Vector-Borne Diseases: Workshop Summary doi:10.17226/21792. National Academies Press (US) Copyright 2016 by the National Academy of Sciences. All rights reserved., Washington (DC).

22. Kraemer MU, Sinka ME, Duda KA, Mylne AQ, Shearer FM, Barker CM, Moore CG, Carvalho RG, Coelho GE, Van Bortel W, Hendrickx G, Schaffner F, Elyazar IR, Teng HJ, Brady OJ, Messina JP, Pigott DM, Scott TW, Smith DL, Wint GR, Golding N, Hay SI. 2015. The global distribution of the arbovirus vectors Aedes aegypti and Ae. albopictus. Elife 4:e08347.

23. Walker T, Jeffries CL, Mansfield KL, Johnson N. 2014. Mosquito cell lines: history, isolation, availability and application to assess the threat of arboviral transmission in the United Kingdom. Parasit Vectors 7:382.

24. Condreay LD, Brown DT. 1986. Exclusion of superinfecting homologous virus by Sindbis virus-infected Aedes albopictus (mosquito) cells. J Virol 58:81–86.

25. Waldock J, Olson KE, Christophides GK. 2012. Anopheles gambiae antiviral immune response to systemic O’nyong-nyong infection. PLoS Negl Trop Dis 6:e1565.

26. Sanchez-Vargas I, Scott JC, Poole-Smith BK, Franz AW, Barbosa-Solomieu V, Wilusz J, Olson KE, Blair CD. 2009. Dengue virus type 2 infections of Aedes aegypti are modulated by the mosquito’s RNA interference pathway. PLoS Pathog 5:e1000299.

27. Morazzani EM, Wiley MR, Murreddu MG, Adelman ZN, Myles KM. 2012. Production of virus-derived ping-pong-dependent piRNA-like small RNAs in the mosquito soma. PLoS Pathog 8:e1002470.

28. Leger P, Lara E, Jagla B, Sismeiro O, Mansuroglu Z, Coppee JY, Bonnefoy E, Bouloy M. 2013. Dicer-2- and Piwi-mediated RNA interference in Rift Valley fever virus-infected mosquito cells. J Virol 87:1631–1648.

29. Bryant B, Macdonald W, Raikhel AS. 2010. microRNA miR-275 is indispensable for blood digestion and egg development in the mosquito Aedes aegypti. Proc Natl Acad Sci U S A 107:22391–22398.

30. Lucas KJ, Roy S, Ha J, Gervaise AL, Kokoza VA, Raikhel AS. 2015. MicroRNA-8 targets the Wingless signaling pathway in the female mosquito fat body to regulate reproductive processes. Proc Natl Acad Sci U S A 112:1440–1445.

31. Liu S, Lucas KJ, Roy S, Ha J, Raikhel AS. 2014. Mosquito-specific microRNA-1174 targets serine hydroxymethyltransferase to control key functions in the gut. Proc Natl Acad Sci U S A 111:14460–14465.

32. Vagin VV, Sigova A, Li C, Seitz H, Gvozdev V, Zamore PD. 2006. A distinct small RNA pathway silences selfish genetic elements in the germline. Science 313:320–324.

33. Luteijn MJ, Ketting RF. 2013. PIWI-interacting RNAs: from generation to transgenerational epigenetics. Nat Rev Genet 14:523–534.

34. Miesen P, Joosten J, van Rij RP. 2016. PIWIs Go Viral: Arbovirus-Derived piRNAs in Vector Mosquitoes. PLoS Pathog 12:e1006017.

35. Girardi E, Miesen P, Pennings B, Frangeul L, Saleh MC, van Rij RP. 2017. Histone-derived piRNA biogenesis depends on the ping-pong partners Piwi5 and Ago3 in Aedes aegypti. Nucleic Acids Res doi:10.1093/nar/gkw1368.

36. Parrish NF, Fujino K, Shiromoto Y, Iwasaki YW, Ha H, Xing J, Makino A, Kuramochi-Miyagawa S, Nakano T, Siomi H, Honda T, Tomonaga K. 2015. piRNAs derived from ancient viral processed pseudogenes as transgenerational sequence-specific immune memory in mammals. RNA 21:1691–1703.

37. Zhang P, Kang JY, Gou LT, Wang J, Xue Y, Skogerboe G, Dai P, Huang DW, Chen R, Fu XD, Liu MF, He S. 2015. MIWI and piRNA-mediated cleavage of messenger RNAs in mouse testes. Cell Res 25:193–207.

38. Goh WS, Falciatori I, Tam OH, Burgess R, Meikar O, Kotaja N, Hammell M, Hannon GJ. 2015. piRNA-directed cleavage of meiotic transcripts regulates spermatogenesis. Genes Dev 29:1032–1044.

39. Post C, Clark JP, Sytnikova YA, Chirn GW, Lau NC. 2014. The capacity of target silencing by Drosophila PIWI and piRNAs. Rna 20:1977–1986.

40. Roiz D, Vazquez A, Seco MP, Tenorio A, Rizzoli A. 2009. Detection of novel insect flavivirus sequences integrated in Aedes albopictus (Diptera: Culicidae) in Northern Italy. Virol J 6:93.

41. Goic B, Vodovar N, Mondotte JA, Monot C, Frangeul L, Blanc H, Gausson V, Vera-Otarola J, Cristofari G, Saleh MC. 2013. RNA-mediated interference and reverse transcription control the persistence of RNA viruses in the insect model Drosophila. Nat Immunol 14:396–403.

42. Chiba S, Kondo H, Tani A, Saisho D, Sakamoto W, Kanematsu S, Suzuki N. 2011. Widespread endogenization of genome sequences of non-retroviral RNA viruses into plant genomes. PLoS Pathog 7:e1002146.

43. Nene V, Wortman JR, Lawson D, Haas B, Kodira C, Tu ZJ, Loftus B, Xi Z, Megy K, Grabherr M, Ren Q, Zdobnov EM, Lobo NF, Campbell KS, Brown SE, Bonaldo MF, Zhu J, Sinkins SP, Hogenkamp DG, Amedeo P, Arensburger P, Atkinson PW, Bidwell S, Biedler J, Birney E, Bruggner RV, Costas J, Coy MR, Crabtree J, Crawford M, Debruyn B, Decaprio D, Eiglmeier K, Eisenstadt E, El-Dorry H, Gelbart WM, Gomes SL, Hammond M, Hannick LI, Hogan JR, Holmes MH, Jaffe D, Johnston JS, Kennedy RC, Koo H, Kravitz S, Kriventseva EV, Kulp D, Labutti K, Lee E, et al. 2007. Genome sequence of Aedes aegypti, a major arbovirus vector. Science 316:1718–1723.

44. Lan Q, Fallon AM. 1990. Small heat shock proteins distinguish between two mosquito species and confirm identity of their cell lines. Am J Trop Med Hyg 43:669–676.

45. Fansiri T, Fontaine A, Diancourt L, Caro V, Thaisomboonsuk B, Richardson JH, Jarman RG, Ponlawat A, Lambrechts L. 2013. Genetic mapping of specific interactions between Aedes aegypti mosquitoes and dengue viruses. PLoS Genet 9:e1003621.

46. Caron M, Grard G, Paupy C, Mombo IM, Bikie Bi Nso B, Kassa Kassa FR, Nkoghe D, Leroy EM. 2013. First evidence of simultaneous circulation of three different dengue virus serotypes in Africa. PLoS One 8:e78030.

47. Fontaine A, Jiolle D, Moltini-Conclois I, Lequime S, Lambrechts L. 2016. Excretion of dengue virus RNA by Aedes aegypti allows non-destructive monitoring of viral dissemination in individual mosquitoes. Sci Rep 6:24885.

48. Kearse M, Moir R, Wilson A, Stones-Havas S, Cheung M, Sturrock S, Buxton S, Cooper A, Markowitz S, Duran C, Thierer T, Ashton B, Meintjes P, Drummond A. 2012. Geneious Basic: an integrated and extendable desktop software platform for the organization and analysis of sequence data. Bioinformatics 28:1647–1649.

49. Guindon S, Dufayard JF, Lefort V, Anisimova M, Hordijk W, Gascuel O. 2010. New algorithms and methods to estimate maximum-likelihood phylogenies: assessing the performance of PhyML 3.0. Syst Biol 59:307–321.

